# Real-Time Visualization of G2L4 Reverse Transcriptase in DNA Repair via Microhomology-Mediated End Joining

**DOI:** 10.64898/2026.03.13.711756

**Authors:** Pangmiaomiao Zhang, Mo Guo, Y. Jessie Zhang, Alan M. Lambowitz, Yi-Chih Lin

## Abstract

Double-strand break repair (DSBR) is essential for genome integrity, yet mechanistic details of error-prone microhomology-mediated end joining (MMEJ) remain unclear. A bacterial group II intron-like reverse transcriptase, G2L4 RT, has been implicated in MMEJ, but how it executes DSBR is unknown. Using high-speed atomic force microscopy (HS-AFM), we directly visualize G2L4 RT-mediated DSBR via MMEJ. We observe that G2L4 RT dimers exhibit RT3a plug protrusion upon DNA engagement and catalyze MMEJ by binding and stabilizing a 4-bp annealed microhomology and filling adjacent single-strand gaps with dNTPs. We also observe Mn^2+^-stimulated terminal transferase activity that generates elongated and branched DNA intermediates prior to ligation. With T4 DNA ligase, we visualize binding near nick sites and real-time nick sealing, which stabilizes the repaired products and suppresses off-pathway branching. These results reveal how G2L4 RT and ligase activities shape MMEJ intermediates and outcomes.

## Introduction

Double-strand breaks (DSBs) arise from exogenous DNA damage sources, such as chemotherapeutics, ionizing radiation and reactive oxygen species.^1^ Even without environmental stress, DNA remains vulnerable during replication, and unrepaired damage can lead to genome instability.^1,2^ In general, DNA damage blocks accurate replication and transcription, drives mutagenesis, and fosters cellular aging.^3^ It also contributes to numerous diseases, most notably cancer, where defective double-strand break repair (DSBR) generates therapy-resistant tumor cells. Therefore, understanding the mechanism of DSBR is critical not only for chromosome biology but also for genome engineering and therapeutic applications.^1^

In mammalian cells, most DSBs are repaired by either homologous recombination (HR) or classical non-homologous end joining (cNHEJ).^4^ HR uses an identical, homologous DNA sequence from the sister chromatid as a template and is therefore considered a high-fidelity DSBR mechanism.^2^ In contrast, cNHEJ is efficient but error-prone, directly rejoining broken DNA ends in a manner that frequently introduces nucleotide deletions and/or insertions at the repair junctions.^1,2,4^ When HR or cNHEJ is compromised, cells can invoke alternative and intrinsically mutagenic pathways, such as microhomology-mediated end joining (MMEJ) and single-strand annealing (SSA).^1,2,5^ These pathways can be distinguished by the length of homologous regions formed on the DNA substrate and by the enzymatic players acting at different stages of repair.^5^

MMEJ initiates after limited 5’-3’ end resection of the DSB, producing short single-stranded (ss) DNA overhangs that contain microhomology segments, typically 5-20 nucleotides (nts) in length, which anneal to each other.^2^ The full MMEJ proceeds through several sequential steps: (1) DNA end recognition, (2) end pairing and microhomology base pairing, (3) removal of non-homologous tails by nucleases, (4) ss-gap filling by DNA polymerases, and (5) ligation by DNA ligases to restore backbone continuity.^6^ Because MMEJ is an error-prone DSBR pathway, the repaired DNA frequently accumulates mutations, including deletions, insertions, and base substitutions.^2,6,7^ In eukaryotes, these repair steps are primarily carried out by specialized polymerases that enable low-fidelity gap filling during MMEJ.^8^

DNA polymerase θ (Pol θ), a promising chemotherapeutic target that is highly expressed in breast and ovarian cancer cells,^9^ is a major mediator of DSBR via MMEJ in eukaryotes.^6,9-13^ Recent studies showed that a *Pseudomonas aeruginosa* chromosomally encoded group II intron-like reverse transcriptase (G2L4 RT) performs DSBR through mechanisms similar to those of Pol θ and possesses multiple enzymatic activities that make it a robust DNA repair enzyme, including the ability to read through DNA lesions, Mn^2+^-stimulated terminal transferase activity, and DSBR via MMEJ.^14,15^ A bacterial group II intron-encoded RT (GsI-IIC RT) was found to have a similar basal DSBR activity, and the DNA repair function of both enzymes rely on conserved structural features of non-LTR-retroelement RTs, a family of enzymes that also includes human LINE-1 and other eukaryotic non-LTR-retrotransposon RTs.^14^ Collectively, these findings suggested that DSBR may be a broadly conserved function of related RTs across diverse organisms.

Mechanistic insights into DSBR have traditionally come from four complementary approaches: (1) in vitro biochemical assays that detect and quantify repair products, (2) in vivo genetic studies that measure the extent and kinetics of DSBR^16^, (3) high-resolution structures of enzyme-DNA complexes resolved by X-ray crystallography or cryogenic electron microscopy (cryo-EM), and (4) single-molecule fluorescence methods that track the kinetics of enzyme-DNA interactions during repair^16,17^. Together, these approaches have revealed important molecular architectures and proposed mechanisms for different DSBR pathways. However, as a long-standing limitation of single-molecule studies, the trade-off between spatial and temporal resolution leaves a gap in directly capturing the structural dynamics of repair enzymes as they execute DSBR.

Among various biophysical techniques, high-speed atomic force microscopy (HS-AFM) is unique in its ability to directly visualize biomolecules in action, including the conformational changes of channels and transporters between functional states, the self-assembly of intrinsically disordered proteins, and protein-DNA interactions under near-physiological conditions.^18^ Previous HS-AFM studies have successfully captured the real-time dynamics of CRISPR-Cas9 during dsDNA cleavage and histone H2A during DNA wrapping.^19-21^ In this study, we employed HS-AFM to directly study the functional activities of G2L4 RT on a designed MMEJ sample. We first characterized the structural dynamics of G2L4 RT, and then tracked its engagement with MMEJ substrates to drive real-time repair. Beyond MMEJ repair, we observed the formation of complex linear and branched products generated by the combined enzymatic activities of G2L4 RT. Finally, we visualized completion of the repair process by capturing T4 DNA ligase activity and the resulting changes in DNA topology. Together, these observations provide a dynamic, mechanistic view of G2L4 RT-mediated DSBR via MMEJ and uncover pathways that may underlie diverse DSBR outcomes.

## Results

### G2L4 RT dimers can dissociate into two monomers in the presence of Mn^2+^

G2L4 RT apoenzyme (APO) forms a homodimer in which each monomeric structure is composed of three hand-like regions, denoted fingers, palm, and thumb (**Fig.1A**, PDB: 9D5X).^15^ A unique insertion of a double helical bundle, denoted the RT3a knot (**Fig.1A**, yellow highlighted), behaves like a plug to block the active site of G2L4 RT.^15^ When G2L4 RT is functionally activated for DSBR, the RT3a plug is extruded, exposing the active site to bind catalytic divalent cations and DNA substrates and function in DNA repair.^15^ In our previous studies, biochemical control of Mn^2+^ ions enhanced G2L4 RT DSBR activities, presumably by weakening the interaction between the RT3a plug and the active site to promote functional activation.^14,15^ However, the mechanism by which Mn^2+^ modulates conformational change and enhances polymerase activity remains unclear at the single molecule level.

**Figure 1.**
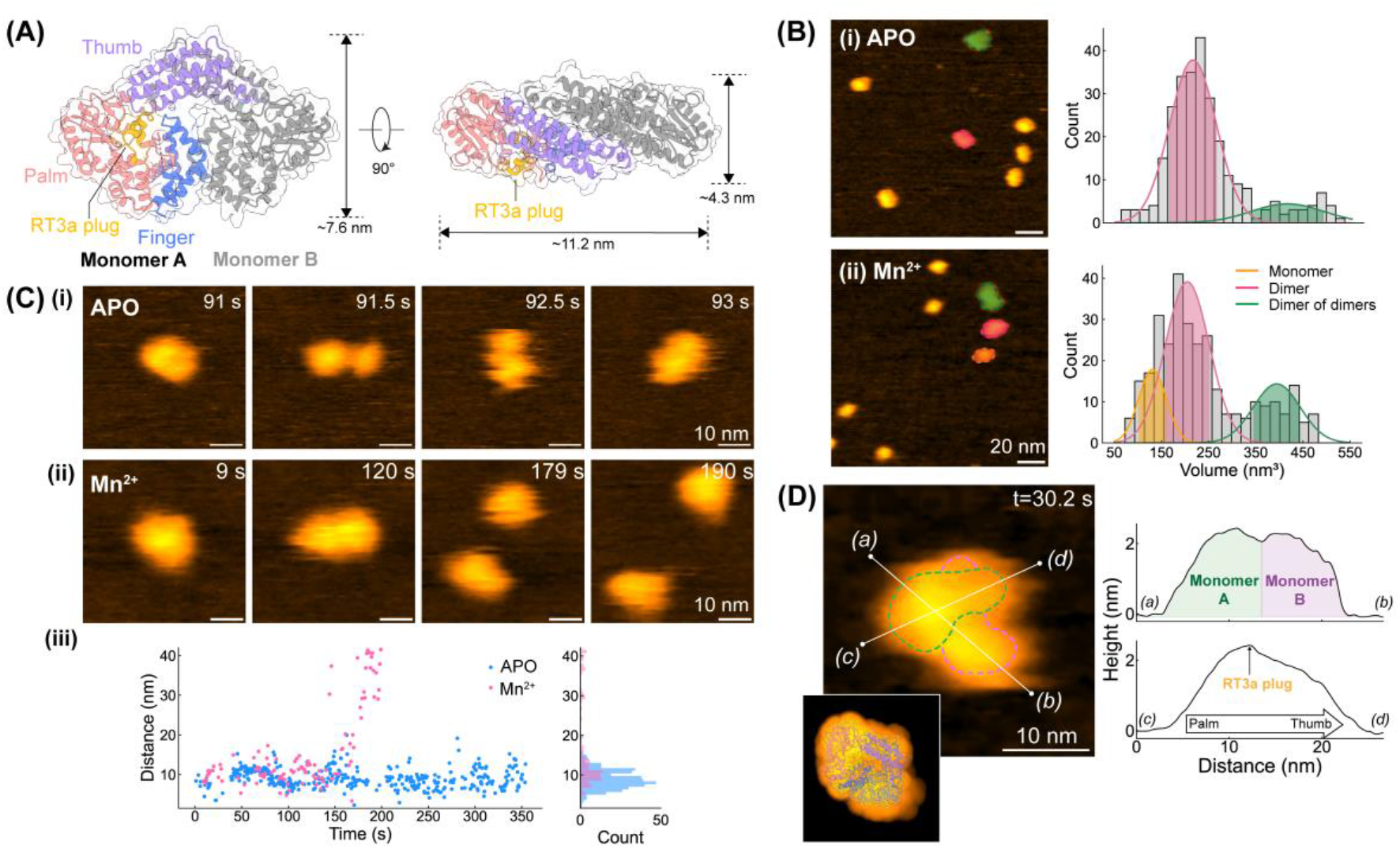
Structural dynamics of G2L4 RT. **(A)** X-ray crystallography structure of the G2L4 RT dimer (PDB: 9D5X; APO state). Three hand-like regions (finger, palm, thumb) and the RT3a plug are highlighted in different colors in Monomer A. **(B)** Representative HS-AFM images of G2L4 RT molecules observed in (i) APO and (ii) Mn^2+^ conditions. G2L4 RT monomers (orange), dimers (pink), and dimer of dimers (green) are assigned based on the particle dimensions and volume. *Right:* Volume histograms from APO (n = 276; two Gaussian fits centered at 214.5 ± 49.4 nm^3^ and 413.7 ± 76.0 nm^3^) and Mn^2+^ conditions (n = 308; three Gaussian fits centered at 131.0 ± 29.4 nm^3^, 204.9 ± 46.9 nm^3^, and 394.6 ± 49.7 nm^3^). **(C)** Time-lapse HS-AFM images of individual G2L4 RT dimer observed in (i) APO and Mn^2+^ conditions. (iii) Tracked monomer-monomer distance over time with a cumulative histogram shown on the right. **(D)** High-resolution HS-AFM image of G2L4 RT dimer (**Fig. S1**, t = 30.2 s) showing the relative positions of two monomers from *(a)* to *(b)*. The height profile from *(c)* to *(d)* corresponds to palm-to-thumb features of a G2L4 RT monomer, with a small height extrusion consistent with the RT3a plug. *Bottom left inset:* simulated AFM topography of G2L4 RT dimer overlaid with the PDB model, showing the best structural match to the observed HS-AFM features.

To characterize the structures of G2L4 RT at different functional states, we first performed HS-AFM experiments to visualize the G2L4 RT particles in either APO or Mn^2+^-activated conditions (**Fig. 1B-1D**, and **Fig. S1**). APO-G2L4 RT particles deposited on mica substrate can be classified as dimers and “dimer of dimers” based on their observed dimensions. This classification is supported by volume analysis, which revealed two distinct particle populations with mean volumes centered at 214.5 nm^3^ and 413.7 nm^3^ (**Fig. 1B**-(i), *n=276*), corresponding to a G2L4 RT dimers and tetramers, respectively. The average volume of the G2L4 RT dimer measured by HS-AFM is larger than that calculated from the corresponding PDB structure (**Fig. 1A**, ∼103 nm^3^). Such volume enlargement is commonly observed in HS-AFM measurements using a sharp tip (∼1 nm radius at apex) and arises mainly from the dynamic nature of proteins, as well as minor tip-sample convolution effects^22^ that widen the apparent surface topography.

Rather than static snapshots, time-lapse HS-AFM movie frames further show that the APO-G2L4 RT dimer exhibits conformational dynamics while preserving its dimeric form (**Fig. 1C**-(i), and **Supplementary Movie S1**). Throughout the observation window, the APO-G2L4 RT dimer maintains the inter-monomer separation at ∼9.2±2.5 nm without dissociation over time (**Fig. 1C**-(iii), blue). The high-resolution HS-AFM image of another APO-G2L4 dimer further shows a bi-lobed particle ∼20 nm in length with ∼2.2 nm extrusions in height (**Fig. 1D, Fig. S1**, and **Supplementary Movie S2**). To interpret these structural features, we simulated the AFM topography of G2L4 RT based on the corresponding PDB structure using an optimized orientation that best matches the HS-AFM data (**Fig. 1D**, bottom left inset). The G2L4 RT dimer observed in HS-AFM images exhibits a distinct small gap separating the finger domains of the two monomers (**Fig. 1D**, line a-b), as well as topological features extending from the palm to the thumb domains with a minor protrusion that likely corresponds to the RT3a plug (**Fig. 1D**, line c-d). Taken together, the bi-lobed morphology and their conformational variability (**Fig. 1C**-(i), **Fig. 1D**, and **Fig. S1**) indicate relative motions of the two palm-finger domains within the dimer. These observations are consistent with previous structural studies showing that G2L4 RT forms dimer with a stable interface at the thumb domains, whereas the palm-finger domains remain more flexible and undergo relative motions.^15^

In the presence of Mn^2+^ ions, HS-AFM images revealed an additional monomer population, in addition to dimers and “dimer of dimers”, with a mean volume centered at 131 nm^3^ (**Fig. 1B**-(ii)). Unlike APO condition, Mn^2+^ enabled dissociation of the G2L4 RT dimer into two monomers (**Fig. 1C**-(ii) and **Supplementary Movie S3**). Molecular trajectories further showed that Mn^2+^-bound dimers display more vigorous relative motions, with an inter-monomer separation in 10.7±2.9 nm prior to dissociation (**Fig. 1C**-(iii)). These observations suggest that Mn^2+^ binding destabilizes the inter-subunit interface present in the APO crystal structure. We hypothesize that this Mn^2+^-induced conformational flexibility may facilitate functional activation by lowering the energetic barrier for RT3a plug displacement and improving substrate access to the active site.

### Structural dynamics of MMEJ DNA substrates with microhomology formation

To investigate the function of G2L4 RT in MMEJ, we designed a DNA substrate with a ss-overhang via annealing a long ssDNA (96 nt, D2) with a short ssDNA (60 nt, D1) (**Fig. 2A**, *Top &* **Table S1**). The ss-overhang of DNA substrate is at the end of D2, terminating in 4-nt complementary sequence CCGG, which can form a 4-bp microhomology (MH) with a second DNA substrate. Here, we refer to the designed DNA substrate as the MMEJ substrate and define the MMEJ dimer as two MMEJ substrates connected by base pairing of the 4 nt microhomology (**Fig. 2D**, *left schematic*). Accordingly, MMEJ dimer would feature two dsDNA ends (60 bp each) connected by a ssDNA-annealed MH-ssDNA regions (32 nt-4 bp-32 nt).

**Figure 2.**
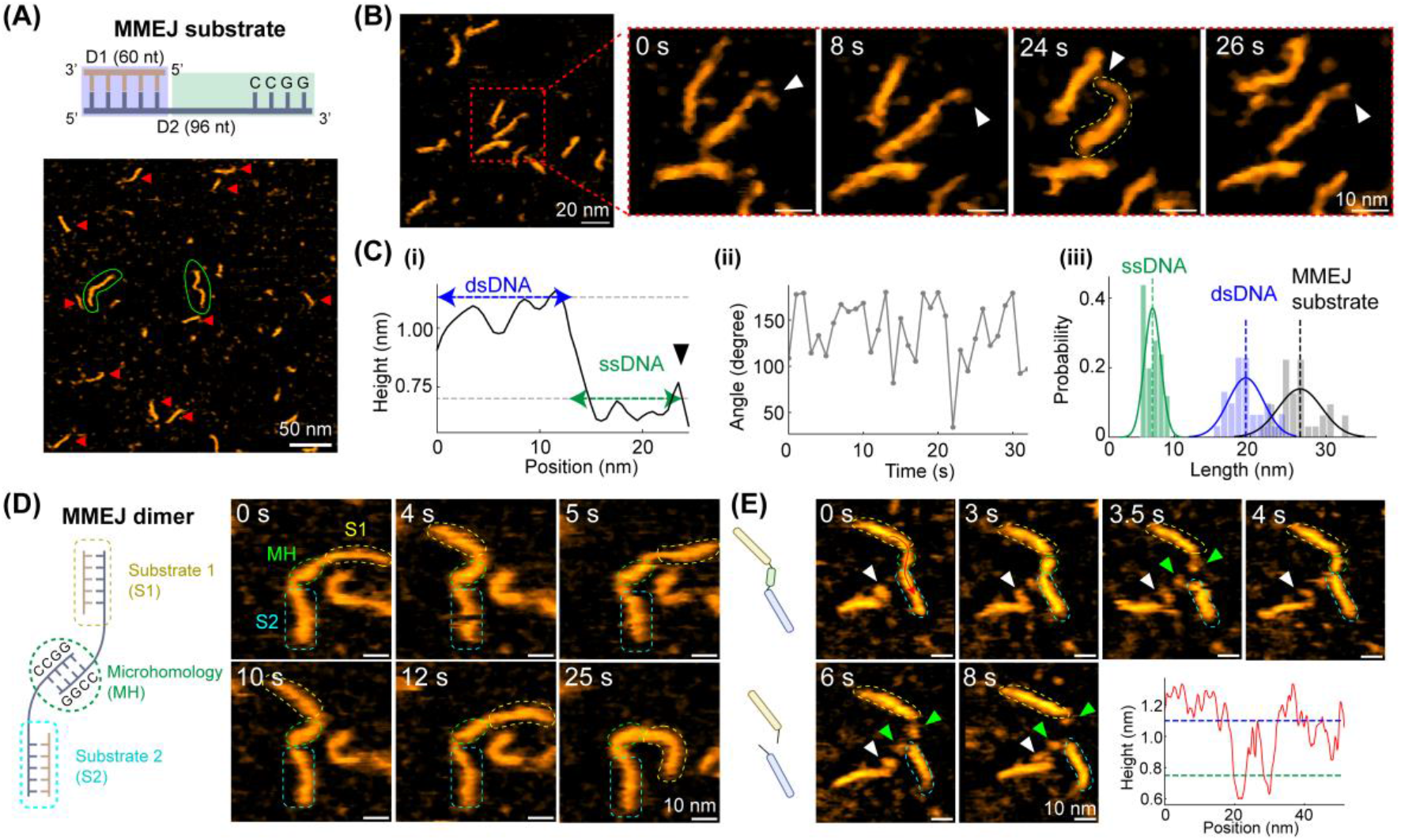
Characterizations of designed MMEJ sample and microhomology (MH) formation. **(A)** Representative HS-AFM image of MMEJ sample deposited on APTES-pretreated mica. Individual MMEJ substrates are marked by red arrows, and MMEJ dimers are indicated by green circles. **(B)** Time-lapse HS-AFM images of a zoomed-in MMEJ substrate show the dynamic breathing of the ssDNA overhang as indicated by white arrowheads. **(C)** (i) Height profile along the backbone of MMEJ substrate at 24 s in **(B)**. Dashed lines represent for the average height of dsDNA and ssDNA, respectively. (ii) Tracked angle between the dsDNA and ssDNA region for the center MMEJ substrate in **(B)**. The length histogram of analyzed MMEJ DNA substrates (n = 32), with an average length of ∼26.6 ± 2.8 nm for full MMEJ substrate, ∼19.4 ± 2.3 nm for dsDNA region, and ∼7.1 ± 1.1 nm for ssDNA overhang, respectively. **(D)** Representative HS-AFM images of a MMEJ dimer with a MH bridge. Two dsDNA ends corresponding to distinct MMEJ substrates (S1 and S2) and MH are labeled based on the height difference and bending geometry. **(E)** Real-time observation of MMEJ microhomology formation and dissociation, indicated by green circles and arrowheads. *Bottom right:* Height cross-section taken along the MMEJ dimer backbone at 0 s.

Based on the previous reports, the B-form DNA has a length of approximately 0.34 nm per bp^23^ and a height of ∼1.5 nm^24^. In contrast, ssDNA are more variable due to its flexibility and depend strongly on factors such as sequence, conformation, and environmental conditions. Empirically, extended ssDNA has a length of ∼0.32-0.68 nm per nt^25,26^, depending on measurement method, with a height approximately half that of dsDNA.^25^ Assuming the ssDNA regions are fully extended, the expected lengths of each dsDNA end and the ssDNA-MH-ssDNA motif are 20.4 nm and ∼21.8∼44.9 nm, respectively. The height contrast between dsDNA and ssDNA provides the basis for distinguishing different regions along the DNA molecules observed in HS-AFM images.

Herein, we performed HS-AFM experiments to study the prepared MMEJ sample in buffer containing 10 mM MgCl_2_, under conditions where MMEJ dimers can form via MH-mediated end joining. The zoomed-out HS-AFM image shows that both MMEJ substrate and MMEJ dimer coexist (**Fig. 2A**), but MMEJ dimers are rare (∼4.5%, **Fig. S2**). Zoomed-in, time-lapse images of a single MMEJ substrate further highlight the height variations along the backbone of MMEJ substrate, where the dsDNA and ssDNA regions are more recognizable based on height difference of ∼0.5 nm (**Fig. 2B, Fig. 2C-i**, and **Supplementary Movie S4**). Statistically, the average measured height of ssDNA and dsDNA are 0.7 ± 0.2 nm and 1.2 ± 0.2 nm (**Fig. S3**), respectively.

In addition, the ssDNA overhang, corresponding to the 3’ end of D2, is highly dynamic with a breathing-like tether motion on substrate (**Fig. 2B**, white arrows). To quantify ssDNA dynamics, we measured the angle formed between dsDNA and ssDNA regions (**Fig. 2C-ii**). The resulting angle distribution exhibits significant fluctuation, ranging from 30° to 180°. It is well-known that ssDNA is much less stiff than dsDNA (persistence length ∼1-6 nm versus ∼50 nm) ^26-28^, which is consistent with the large bending fluctuations observed here.

We also quantified the length distributions of MMEJ substrates that show the average length of 19.4 nm for the dsDNA region, 7.1 nm for the ssDNA overhang, and a total contour length of ∼26.6 nm for MMEJ substrates (**Fig. 2C-iii**). The length of dsDNA region is comparable to the expected dimension; however, the ssDNA region is much shorter than the predicted length. Based on our data, the length of one nucleotide within the designed ssDNA region is approximate 0.2 nm, suggesting that the ssDNA is in a more folded geometry than extended conformations. We suspect the Mg^2+^ in buffer can partially neutralize highly negatively charged phosphate backbone of ssDNA and reduce the electrostatic repulsion between different parts of the same strand.^23^

In addition to individual MMEJ substrates, we observed some MMEJ dimers with an annealed MH-region at their centers (**Fig. 2A**, green circles) in the absence of bound protein. The MMEJ dimers have two dsDNA ends and behave like free-moving handles, whereas the MH-region and certain ssDNA gaps act like a hinge (**Fig. 2D** and **Supplementary Movie S5**). This bending geometry is associated with the distinct mechanical properties between dsDNA and ssDNA. dsDNA is relatively rigid and linear-like, whereas ssDNA is highly flexible and easily bent. Moreover, we captured the association and dissociation of MH region (**Fig. 2E** and **Supplementary Movie S6**). Specifically, the MH region is formed by contacts between the two ss-3’-overhangs of MMEJ substrates at the beginning, where the height profile along the MMEJ dimer backbone clearly shows the height variations across two ssDNA gaps (∼0.8 nm in height) and the central MH region (∼1.1 nm in height) (**Fig. 2E**, bottom right plot). After 6.0 s, the MH dissociates, producing two freely moving ss-overhangs on distinct MMEJ substrates that slightly diffuse apart at the end. Combining the low MMEJ dimer population and observed MH dissociation, the 4 nt microhomology-mediated end joining is dynamic and reversible under our imaging conditions.

Since DSBR via MMEJ requires the recruitment of dNTPs to fill ss-gaps, we also investigated the structural features of MMEJ dimer preincubated with 1 mM dNTPs for 3 hours prior to HS-AFM imaging. Supplementation with dNTPs results in a higher fraction of MMEJ dimers observed, ∼17.4% of total analyzed MMEJ substrates (**Fig. S2** and **S4**). This is ∼3.9-fold higher than that observed without dNTPs. Notably, the MMEJ dimers under this condition are stable, with no dissociation observed, and exhibit sharp bending geometry, indicating DNA backbone remains unrepaired at ssDNA regions (**Fig. S4**). Furthermore, the ss-gaps are not clearly distinguishable along the backbone of MMEJ dimer based on height difference, and thus we expect that dNTPs bind to the ssDNA gaps. Altogether, these single-component observations establish a baseline for probing the functional dynamics of G2L4 RT during DSBR via MMEJ.

### Functional Activities of G2L4 RT on MMEJ sample

Biochemical and structural studies have previously proposed a molecular mechanism of G2L4 RT-mediated DSBR via MMEJ in the presence of Mn^2+^, generally including that this reverse transcriptase binds to the ssDNA overhang of MMEJ substrate, promotes MH-mediated end joining formation, and then subsequently seals the ss-gaps that bind complementary dNTPs.^14,15^ Although these studies provide great structural insights, it is unclear how G2L4 RT mediates DSBR via MMEJ at the single-molecule level. Here, we adopted a stepwise experimental strategy to visualize the G2L4 RT binding and G2L4 RT-mediated DSBR on the designed MMEJ sample under distinct biochemical controls (**Table S2**).

To investigate the molecular interactions between G2L4 RT and the MMEJ sample, we preincubated both samples in buffer containing 10 mM Mg^2+^ at room temperature prior to HS-AFM imaging. Based on HS-AFM results, we frequently observed protein-DNA complexes (**Fig. 3A**), including (i) a G2L4 RT dimer binding to one MMEJ substrate in the ssDNA region, (ii) a G2L4 RT dimer clamping the MH region, bridging two MMEJ substrates, and (iii) G2L4 RT dimer binding to the terminal dsDNA regions of a MMEJ dimer substrate. Among these protein-DNA complexes, G2L4 RT binding to the 3’-end of a ssDNA overhang (**Fig. 3A**-(i)) was the most frequently observed, consistent with the previously proposed MMEJ mechanism.^14,15^ Interestingly, the MMEJ dimer clamped by a G2L4 RT dimer does not dissociate (**Fig. S5 & Supplementary Movie S7**), indicating that G2L4 RT binding can promote and stabilize MH-mediated end joining formation. This is also supported by the observation that ∼24.6% of the DNA molecules are present as MMEJ dimers under this condition **(Fig. S2)**.

**Figure 3.**
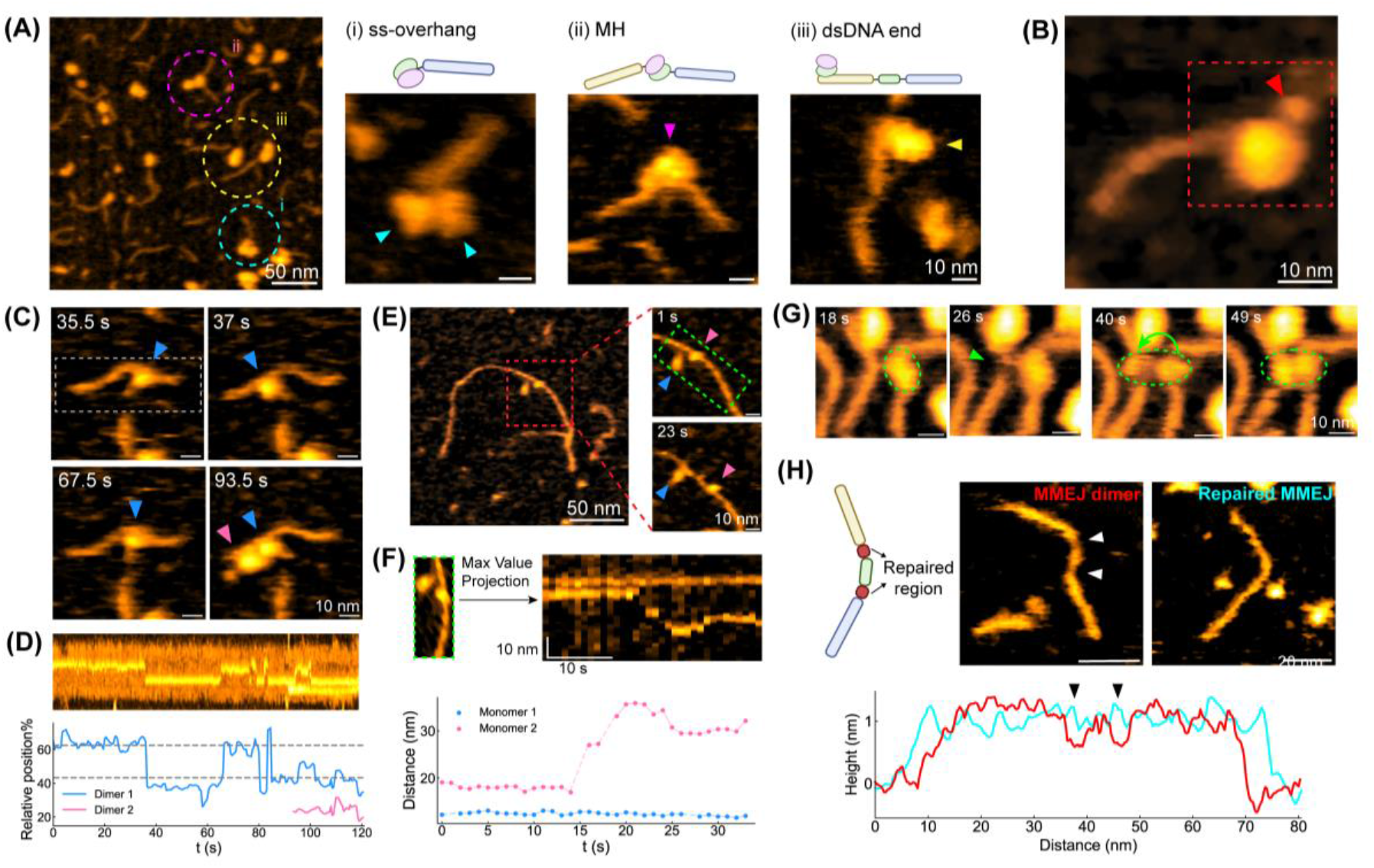
Functional activities of G2L4 RT molecules on the designed MMEJ sample. **(A)** Representative HS-AFM images of the G2L4 RT dimer binding to MMEJ samples at the regions of (i) ss-overhang of MMEJ substrate, (ii) MH and (iii) dsDNA end of the MMEJ dimer in the presence of 10 mM Mg^2+^. **(B)** Representative G2L4 RT dimer exhibits a structural protrusion of ∼3 nm (red arrow) after binding to MMEJ substrate, which is likely the RT3a plug released from its structure. **(C)** Real-time observations of G2L4 RT dimer oscillate between two ss-gaps of a MMEJ dimer, where are supposedly to be repaired by dNTPs and G2L4 RT in the presence of Mn^2+^ cations. **(D)** Kymograph of the G2L4 RT’s position along the backbone of MMEJ dimer, where the dashed lines indicate the relative positions of two ss-gaps in MMEJ dimer. **(E-F)** G2L4 RT monomer slides along the DNA substrate. **(G)** A G2L4 RT dimer binding to MH center of one MMEJ dimer exhibits a conformational change to allow one of the monomeric G2L4 RT hoping to MH center of another MMEJ dimer. **(H)** Representative image of MMEJ dimer before and after G2L4 RT repair with dNTPs. The height profiles along the MMEJ dimers highlights a height difference ∼ 0.4 nm at two ss-gaps, pointed by arrows.

When G2L4 RT dimer bound to MMEJ substrate, we commonly found a small, flexible protrusion next to the observed protein, ∼2 nm in height (**Fig. 3B, Fig. S6**, and **Supplementary Movie S8**). This topological protrusion aligns with the extruded RT3a plug when G2L4 RT in an activated conformation binds to a snapback DNA substrate (**Fig. S6B**; PDB: 9D4S).^15^ Time-lapse HS-AFM images show that when G2L4 RT was not bound to a MMEJ substrate (**Fig. S6A**, after t = 19.5 s), the RT3a plug was still free-moving, enabling a search for an acceptable substrate. In addition to the single protrusion, we also found two RT3a plug protrusions from both monomers of a G2L4 RT dimer bound or not bound to a MMEJ substrate **(Fig. S7)**. These observations support the conformational change of G2L4 RT dimer with RT3a plug release when it engages a DNA substrate and adopts an active configuration for DSBR. This molecular feature is consistent with previous models proposing asymmetric activation within the dimer, in which a leading monomer adopts an active state through RT3a plug extrusion whereas the trailing monomer remains inactive.

Next, we evaluated the G2L4 RT function in repairing the MMEJ sample by supplementing 1 mM dNTPs and 1 mM Mn^2+^ in liquid cell to initiate its DNA repair activities. The divalent cations coordinate with conserved aspartate residues in the G2L4 RT active site, stabilizing both the DNA substrate and the incoming dNTPs for repair.^14,15^ After addition of dNTPs and Mn^2+^ for one hour, both repaired and unrepaired MMEJ dimers, as well as G2L4 RT monomers and dimers, were observed **(Fig. S8)**. Since G2L4 RT can repair the MMEJ dimer, we further investigated how G2L4 RT monomers or dimers perform DSBR via MMEJ mechanism. Notably, a G2L4 dimer (dimer 1) hopped between two strong binding sites, which correspond to the positions of ss-gaps along a MMEJ dimer (**Fig. 3C-D & Supplementary Movie S9**). Another G2L4 dimer (dimer 2) entered the field of view and then bound to the dsDNA end of the MMEJ dimer away from the MH at ∼94 s.

We constructed a kymograph to track the positions of G2L4 RT dimers in this HS-AFM move by projecting the maximum height along the y-axis of each frame (**Fig. 3C**, analyzed area within white dashed rectangle as shown at 35.5 s) as a single vertical line (**Fig. 3D**, top). The resulting kymograph clearly shows that repositioning of the G2L4 RT dimer matches the ss-gap sites that it would repair (**Fig. 3D**). Both dimers display single RT3a protrusion, indicating they are in the active state. Our results suggest that G2L4 RT dimers can repair dsDNA by repositioning at ss-gaps on both sides of the MH region.

We did not observe a G2L4 RT dimer dissociating into two monomers while bound to MMEJ samples. However, some G2L4 RT monomers also interact with MMEJ substrates. We observed a G2L4 monomer bridging MH formation in real-time **(Fig. S9)**. In addition, another observed G2L4 monomer bound to a long DNA molecule under conditions where G2L4 RT and MMEJ samples were preincubated with dNTPs and Mn^2+^ for 6 hours (**Fig. 3E-F**). The G2L4 RT monomer moves along dsDNA backbone in a pause-and-slide manner, where the sliding velocity can reach ∼1.9 nm/s from 14 s to 24 s **(Fig. 3F)**. Considering a length of 0.34 nm/bp for dsDNA, the sliding velocity corresponds to approximately 6 bp/s.

In addition, we also observed a real-time conformational change of a G2L4 RT dimer at the center MH of a MMEJ dimer, where one of the monomers hopped to the central MH of a nearby MMEJ dimer (**Fig. 3G**). The G2L4 RT dimer tightly holds two MMEJ dimers by bridging their central MH regions together. The capability of G2L4 RT to rejoin different DNA substrates may result in the formation of complex DNA repair products, such as dsDNA with branched morphology, which will be addressed in the next section (**Fig. 4**).

**Figure 4.**
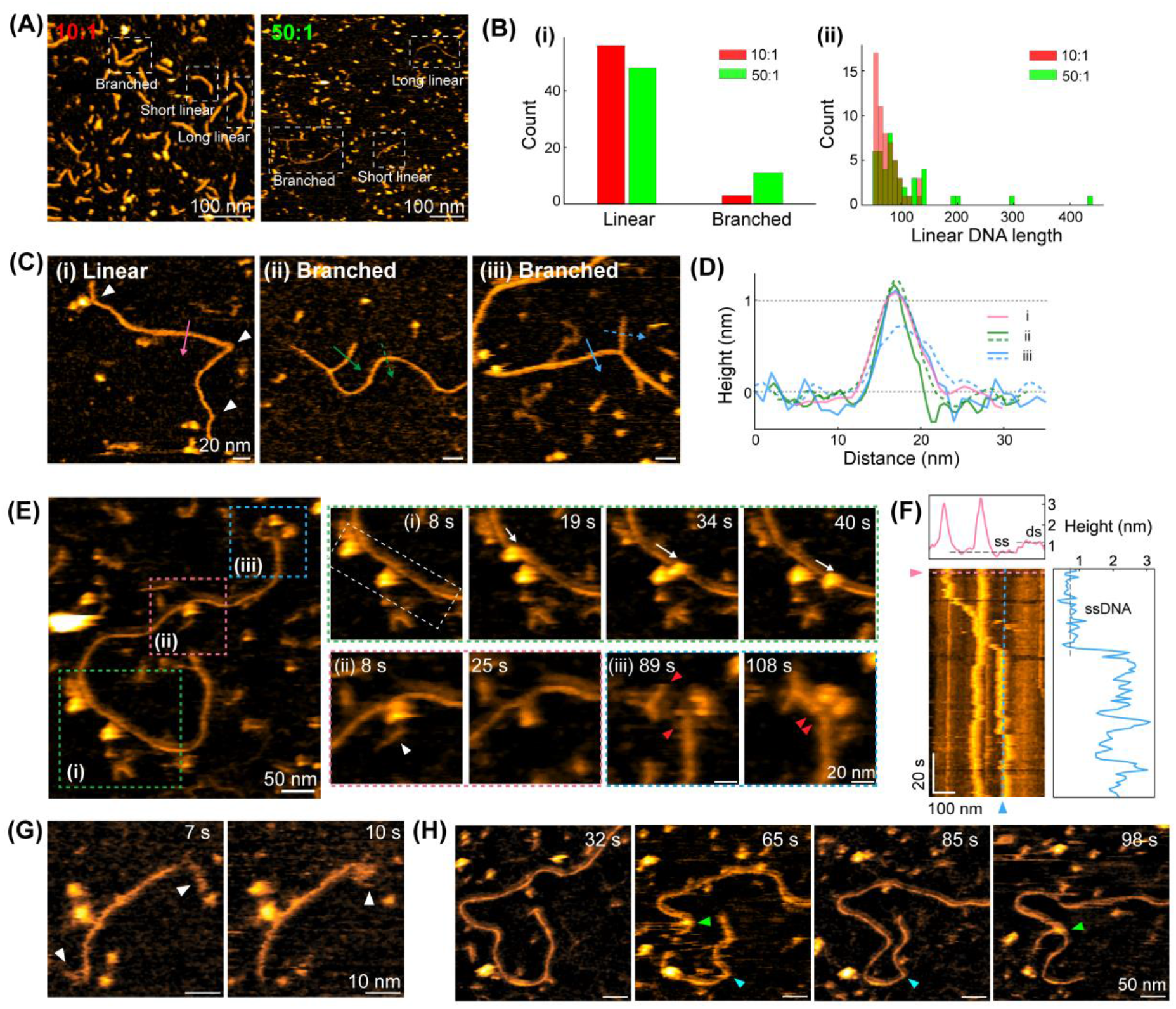
Diverse DNA products after mixing G2L4 RT and MMEJ sample with dNTPs, Mn^2+^ and Mg^2+^ cations for 6 hours, but without T4 ligase. **(A)** Representative HS-AFM images of DNA products produced by G2L4 RT-mediated DSBR under different molar ratio of [G2L4]:[MMEJ]. Based on the observed morphology, different types of DNA molecules are highlighted with white dashed boxes. **(B)** Statistical analysis of the observed DNA products, including (i) DNA morphology distribution and (ii) length histogram of observed linear DNA molecules at [G2L4]:[MMEJ]=10:1 (red) and 50:1 (green) molar ratio, respectively. **(C)** Representative long, linear and branched DNA products via G2L4 RT-mediated DSBR. **(D)** DNA backbone thickness on long, linear and branched DNA products, as shown in **(C). (E)** A representative HS-AFM image of several G2L4 RT molecules interacting with a long, linear DNA molecule. Three colored regions highlight the dynamic events of G2L4 RT dimer (i) sliding on ssDNA, (ii) mediating the formation of ssDNA branch, and (iii) bridging two DNA ends highlighted by red arrowheads. **(F)** Kymograph of the positions of two G2L4 RT dimers on the DNA backbone generated from region of white dashed box in **(E)**-(i). (*Top* and *right*) Maximum height profiles along x (pink dashed line) and y directions (blue dashed line). **(G)** A linear DNA product with ss-overhangs of ∼9 nm and ∼26 nm in length at two ends highlighted by white arrowheads, indicating the terminal transferase activity of G2L4 RT. **(H)** Rearrangement of linear DNA segment from the end (cyan arrow) to the middle of DNA backbone forms a new branch (green arrow).

Except for the complexes of G2L4 RT and DNA substrates, we also compared the MMEJ dimer before and after DSBR by adding G2L4 RT, dNTPs and Mn^2+^ (**Fig. 3H**). Here, the two ss-gaps bridged by the MH region are expected to be filled and converted into dsDNA, with two single gaps (nick sites) remaining for ligation. The repaired MMEJ dimer exhibits a similar length to the unrepaired one, but the original two ss-gaps become thicker, approaching the thickness of dsDNA (**Fig. 3H**). Height profiles along the backbone of unrepaired and repaired MMEJ dimers show an increase in height at the two ssDNA-gaps from ∼0.7 nm to ∼1.0-1.2 nm, which indicates successful G2L4 RT-mediated DSBR via MMEJ mechanism. Consistent with these observations, quantitative height tracking along the DNA backbone over time further shows that MMEJ dimers in the MMEJ-only sample exhibit large height fluctuations and dynamic ss-gap appearance (**Fig. S10A**). In contrast, dNTPs increase dimer stability and reduce gap visibility (**Fig. S10B**), whereas the addition of dNTPs and Mn^2+^ largely eliminates ss-gaps, with only minor height falls occasionally observed (**Fig. S10C**).

Altogether, our observations and analysis reveal that G2L4 RT dimer undergoes conformational activation when it engages on MMEJ substrates and MMEJ dimers. We observed multiple functional activities of G2L4 RT, ranging from interactions with MMEJ substrates/dimers to repairing them through the MMEJ mechanism. Furthermore, we demonstrate that G2L4 RT facilitates DNA repair via the MMEJ mechanism as a dimer in the presence of Mn^2+^. Additionally, Mn^2+^ activation of G2L4 RT favors RT3a plug extrusion that enables DNA binding and supports subsequent functional behaviors, including repositioning and sliding.

### MMEJ is an error-prone DSBR process before resealing nicks via T4 DNA ligase

Based on our MMEJ substrate designs, G2L4 RT is expected to repair the MMEJ dimer in the presence of Mn^2+^ and dNTPs, which produces an expected DNA product with two remaining single-strand breaks (nicks) along the dsDNA backbone before T4 ligation (**Fig. 3H;** cyan). However, following an extended 6-hour preincubation, we commonly observed long DNA products (**Fig. 4A**; >80 nm), in either linear or branched forms, compared to the expected MMEJ dimer product (**Fig. 3H**; ∼60 nm in length). Since MMEJ is an error-prone DSBR process, G2L4 RT may play a critical role in bridging and rejoining different DNA substrates (**Fig. 3G**) before T4 ligase seals the nicked sites on the DNA backbone. Furthermore, the polymerase activity of G2L4 RT can also support terminal transferase activity, which is the template-independent addition of dNTPs to the 3’-hydroxyl end of DNA and is particularly stimulated by the presence of Mn^2+^.^14^ This would also contribute to the formation of unexpected DNA products.

Here, we first tested the impact of the molar ratio of G2L4 RT to the MMEJ sample (10 nM) on the repair of DNA products. When the [G2L4 RT]:[MMEJ] molar ratio is ten-fold, most DNA products are linear with lengths less than 100 nm (**Fig. 4A-B**; red), where both MMEJ substrates and dimers are repaired with a thickness similar to dsDNA. While increasing the molar ratio to 50:1, the lengths of most DNA products are comparable to those observed at the 10:1 molar ratio. However, a few observed DNA molecules are much longer than expected (**Fig. 4B-(ii)**; green, >100 nm). Statistically, we observed more branched DNA at the 50:1 ratio (11 out of 59) than at the 10:1 ratio (3 out of 59) (**Fig. 4B-(i)**). The formation of longer and branched DNA products could be driven by enhanced, Mn^2+^-stimulated terminal transferase activity of G2L4 RT, which extends 3’ ssDNA ends by addition of non-templated nucleotides and commonly generates longer linear DNA products upon repair.^14^ These extended ss-overhangs may increase the probability of forming new MH interactions with other DNA ends, thereby promoting end pairing and rejoining between different DNA substrates.

We closely inspected the fine structures of these unexpected long DNA products, including linear and branched forms (**Fig. 4C**). The long, linear DNA products are mostly repaired as dsDNA with a backbone thickness of ∼1.0-1.2 nm. However, a few highly localized, non-smooth bends (**Fig. 4C-(i)**; white arrow) are frequently observed. These bending positions are the possible nicked sites with a less mechanical strain compared to the unbroken dsDNA. For branched DNA products, the short branches can be either in dsDNA or ssDNA form (**Fig. 4C-(ii), -(iii)** and **Fig. 4D**; green and blue). Our results indicate that following non-templated elongation at 3’ ssDNA ends mediated by G2L4 RT, the newly formed MH sites may locate internally and not necessarily at the 3’-end of ssDNA. Thus, the residual ssDNA region between the new MH site and the 3’ end will become a ssDNA flap at the branch point, which may remain reactive and capable of binding other ssDNA overhangs.

To investigate the function of G2L4 RT in DNA elongation and branch, we characterized the molecular interactions between G2L4 RT and long DNA products (**Fig. 4E** and **Supplementary Movie S10**). Three distinct enzymatic events were found in this representative HS-AFM movie, including (i) G2L4 RT dimer sliding on ssDNA backbone, (ii) formation of an ssDNA flap mediated by G2L4 RT dimer, and (iii) DNA end bridging via a G2L4 RT dimer. We generated a kymograph (same approach as **Fig. 3D**) to analyze G2L4 RT dimer movement and the underlying DNA topology in region (i) (**Fig. 4F**). One G2L4 RT dimer remaining stationary at center, while a second dimer slides continuously for ∼108 nm between 8 s and 40 s, with an average velocity of ∼3.4 nm/s. Cross-sectional height profiles confirm that the G2L4 RT dimer is sliding along ssDNA (**Fig. 4F**, *top* and *right panel*). Although DSBR was not observed, molecular sliding reflects a high affinity of G2L4 RT at ssDNA region and may represent a search mechanism for potential repair sites. In region (ii), a G2L4 RT dimer bound to a junction, where a flap ssDNA attached to the main dsDNA after protein dissociation at 25 s. This flap ssDNA branch likely results from incomplete unidirectional repair.^14^ In region (iii), a G2L4 RT dimer is bridging two dsDNA blunt ends. This phenomenon is consistent with our previous observations regarding G2L4 RT’s affinity for blunt ends (**Fig. 3A**), and its ability to bring non-complementary ends into proximity, similar to other polymerases.^29^ Collectively, these findings reveal G2L4 RT dimer behaviors when encountering complex DNA substrates, highlighting its potential capability to search for repair sites, generate ssDNA flaps and bridge distinct DNA strands.

Furthermore, we observed terminal transferase activity of G2L4 RT molecules. A specific DNA product exhibits two flexible ssDNA overhangs (**Fig. 4G**), one of which is ∼25.5 nm in length, substantially longer than the ssDNA overhang measured in the MMEJ substrate (6.0 ± 2.7 nm, **Fig. 2C**). Beyond just the end products, we also observed dynamic interactions between ssDNA strands and G2L4 RT monomer and dimer species (**Fig. S11**). We not only captured the interaction between a G2L4 RT monomer and a ssDNA overhang (**Fig. S11A**) but also obtained real-time ssDNA elongation from ∼29 nm to ∼35 nm over 92 s by the adjacent G2L4 RT molecules (**Fig. S11B**). These data demonstrate the terminal transferase activity of G2L4 RT, which drives ssDNA extension and the generation of longer DNA products.

Interestingly, one of our HS-AFM movies showed the self-rearrangements of the DNA backbone at a nicked site to form branched morphology (**Fig. 4H** and **Supplementary Movie. S11**). Here, an initially intact linear DNA product dissociated at the nicked site (**Fig. 4H;** cyan arrows at 65 s and 85 s), and then a separated branch rejoined to the middle of the original DNA backbone (**Fig. 4H;** green arrows at 98 s). There was a G2L4 RT molecule transiently bound to this rejoining site at 65 s, suggesting its higher binding affinity for nicked site. These observations indicate that the G2L4 RT-mediated repair products before T4 ligation are structurally unstable and prone to self-rearrangements of the DNA backbone at nicked sites to form branched morphology.

Altogether, our HS-AFM experiments reveal that G2L4-mediated MMEJ repair is an error-prone process yielding a diverse population of DNA products, including longer linear and branched topologies with branches, prior to DNA ligation. The formation of these complex structures is driven by a combination of G2L4 molecular behaviors, including repair site scanning, ssDNA flap formation, DNA end bridging and terminal transferase activity. Furthermore, we identified that the pre-ligation nicks on the DNA backbone introduce intrinsic strand instability that could cause the repair intermediates to undergo structural self-rearrangement.

### G2L4 RT-mediated DSBR via MMEJ Mechanism with T4 DNA ligase

DNA ligation is the final step to complete a DSBR process via MMEJ, which converts damaged DNA structures into stable, intact dsDNA products. This step involves cellular DNA repair enzymes, like DNA ligases activated by ATP, to seal the single-nucleotide breaks (nicks) in the phosphodiester backbone, a reaction that G2L4 RT cannot complete independently.^30^ Here, we used T4 DNA ligase, originated from bacteriophage, which requires ATP and Mg^2+^ to seal DNA nicks.^31^ Structurally, T4 DNA ligase is composed of three structural domains (**Fig 5A**; PDB: 6DT1), each contributing to its ligation activity.^31,32^ The N-terminal DNA-binding domain (DBD) mediates nick recognition and enzyme positioning for catalysis. Oligonucleotide-binding (OB) fold domain together with nucleotidyl-transferase (NTase) domain enables AMP transfer and phosphodiester bond formation to seal the nick.

**Figure. 5.**
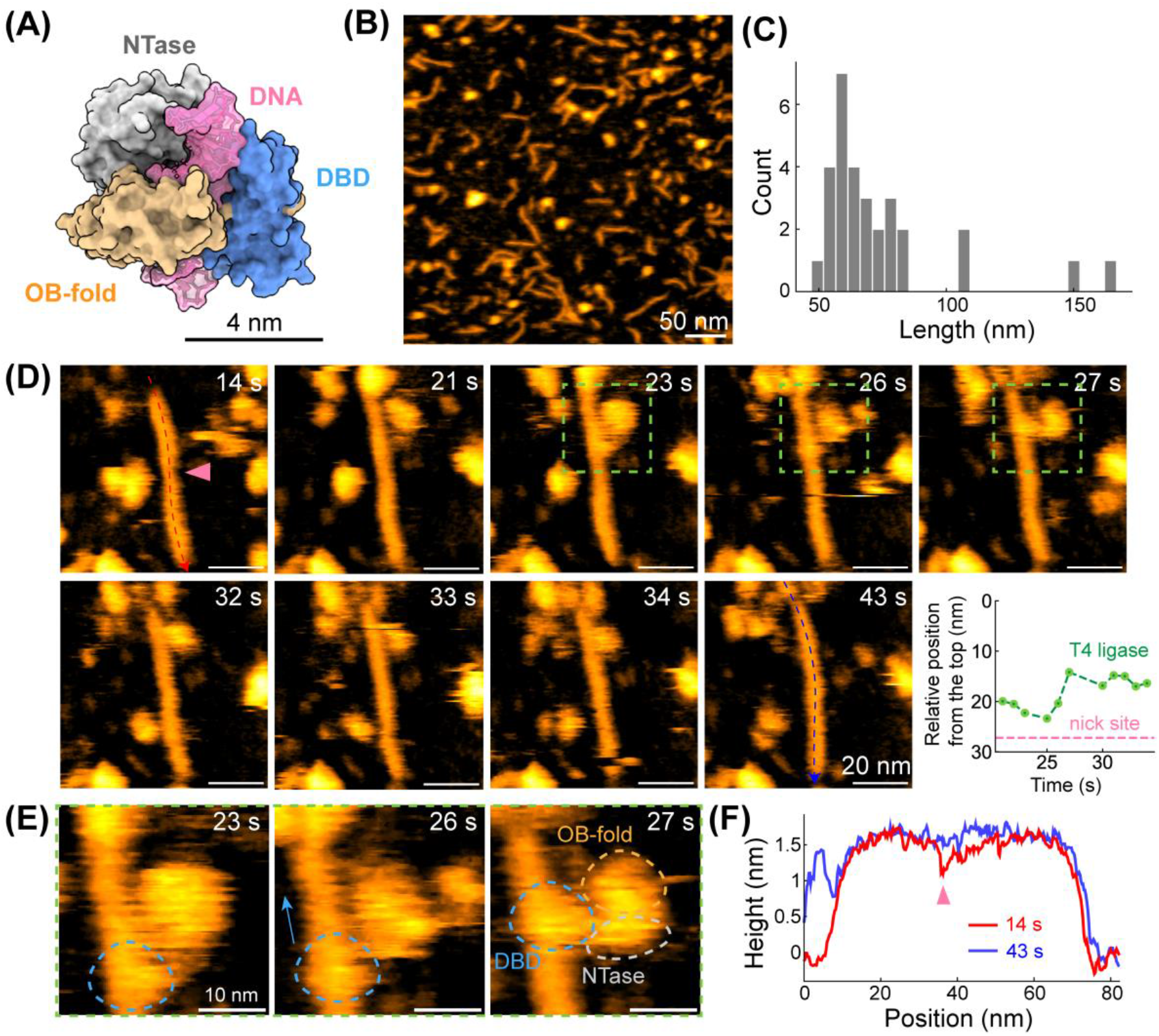
DNA ligation of MMEJ samples via T4 DNA ligase with dNTPs, Mn^2+^, Mg^2+^, and ATP. **(A)** Structure of T4 DNA ligase complexed with DNA intermediate (PDB: 6DT1).**(B)** Representative HS-AFM image of DNA products after DNA ligation. **(C)** Length histogram of DNA products after ligation (n = 30). **(D)** Real-time visualization of T4 DNA ligase interacting with a repaired MMEJ dimer. *Bottom-right:* Relative distance between where the ligase engages and the top of MMEJ dimer. Pink dashed line indicates the position of the nick site observed at 14 s. **(E)** High-resolution images showing domain motion of T4 DNA ligase, zoomed in from **(D)**. Three visible domains of T4 ligase are highlighted by dashed circles. **(F)** Height profile along DNA backbone at t = 14 s and 43 s, which represent to time points before and after DNA ligation.

To first investigate how MMEJ repair product is influenced by the addition of DNA ligase, we pre-incubated T4 DNA ligase (0.04 U) with a mixture of G2L4 RT and the MMEJ sample at a molar ratio of 10:1 in buffer containing 1 mM dNTPs, 10 mM Mg^2+^, 1 mM Mn^2+^, and 1 mM ATP prior to HS-AFM measurements. In the presence of T4 DNA ligase, we observed repaired DNA products that were predominantly MMEJ dimers, with only a few DNA molecules exhibiting lengths (> 100 nm) slightly longer than the designed MMEJ product (**Fig. 5B**). This outcome is different than previous condition containing only G2L4 RT and extended preincubation, where long linear and branched DNA products were frequently observed (**Fig. 4A-4C**).

Notably, we captured the binding of a T4 DNA ligase molecule, which is smaller than G2L4 RT dimer and exhibits a conformation consistent with PDB structure, to a single-nucleotide nick site on a MMEJ dimer (**Fig. 5D-E** and **Supplementary Movie S12**). Before ligase arrival at 14 s, the MMEJ dimer exhibited a distinguishable nick site at ∼27 nm from the top end of DNA, characterized by a ∼0.3 nm lower height (**Fig. 5F**) and narrower width compared with adjacent dsDNA regions. The T4 ligase associated with the DNA backbone between 21 s and 27 s, during which one structural domain appeared to slide along the DNA. The ligase then underwent a conformational rearrangement between 27s and 30 s, followed by the gradual engagement of its core domain with the DNA backbone until 34 s. Analysis of the protein trajectory along the DNA backbone shows that the T4 ligase localized adjacent to the nick site (**Fig. 5D**, *bottom-right panel*). After the ligase dissociated at 43 s, the former nick site was sealed, with the local height increasing to ∼1.5 nm (**Fig. 5F)**. These observations demonstrate that T4 DNA ligase is functionally activated upon binding to nick sites and completes DNA ligation in the presence of ATP.

Moreover, owing to the high spatial and temporal resolution of HS-AFM imaging, we could clearly distinguish three structural domains of T4 DNA ligase during ligation (**Fig. 5E**). The initial contact phase between 21 s and 27 s likely corresponds to scanning behavior mediated by the DBD, which is consistent with its role in nick recognition. Following initial recognition and a conformational transition, the catalytic core of the ligase, comprising the NTase and OB-fold domains, gradually engages the nick site to complete ligation between 30 s and 43 s. Overall, we directly visualized DNA ligation in real-time, representing the crucial final step of the MMEJ pathway that stabilizes repair intermediates and prevents error-prone rearrangements arising from local DNA instability before completion of the DSBR process.

## Discussion

HS-AFM enables the direct visualization of structures, dynamics, and protein-DNA interactions, in contrast to static structural biology or bulk biochemical assays that often miss reaction processes and transient intermediates. Using HS-AFM, we visualized G2L4 RT-mediated DSBR via MMEJ in real-time, thereby bridging the gap between static crystal structures and functional DNA repair mechanisms. Our previous work^14,15^ explored G2L4 RT structure and functions from molecular biology and structural biology perspectives, but the mechanism of how G2L4 RT performs DNA repair remained hypothetical. Here, our HS-AFM data support a more detailed mechanistic model for G2L4 RT-mediated MMEJ repair (**Fig. 6**).

**Figure 6.**
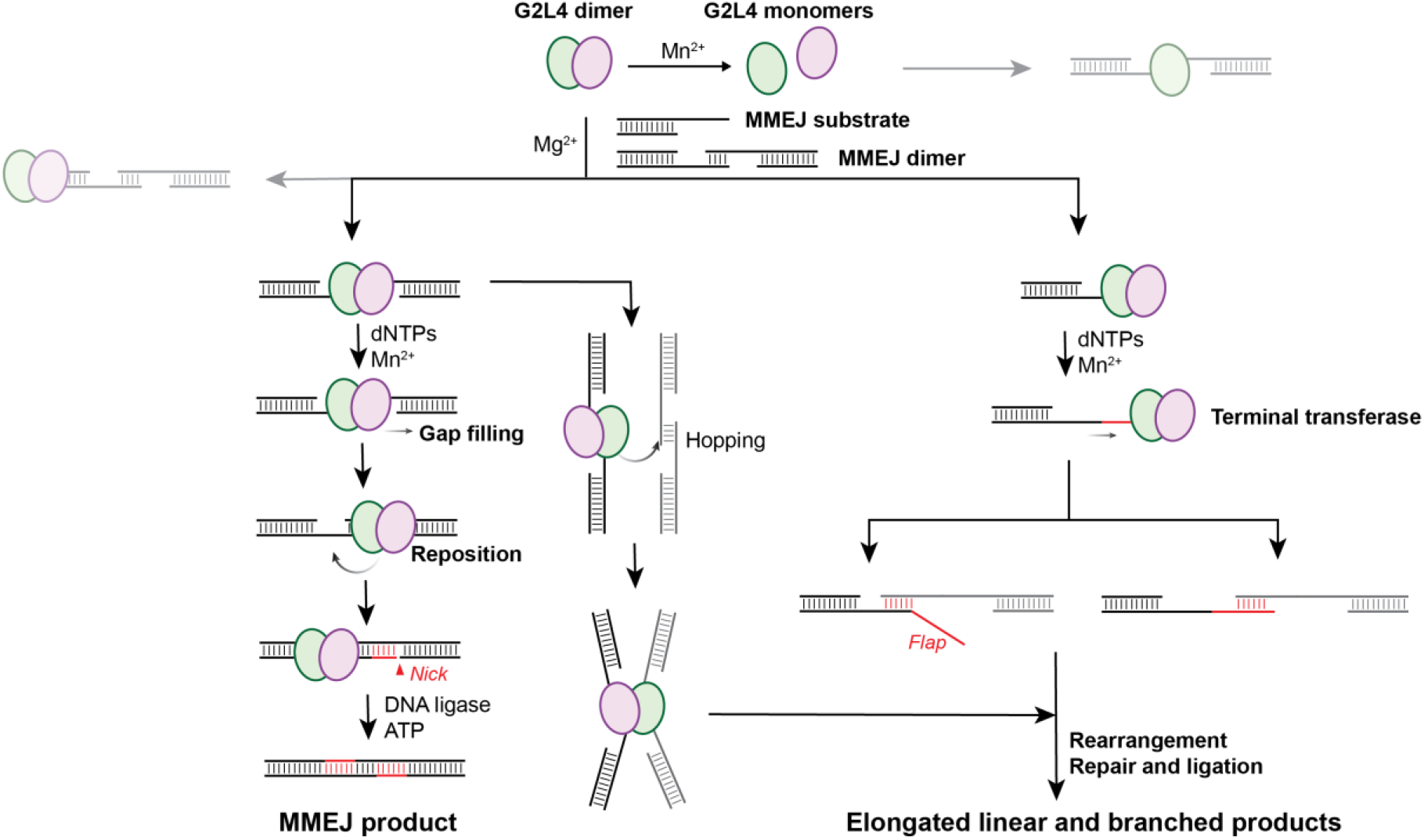
Proposed G2L4 RT-mediated DSBR via MMEJ mechanism. Schematic summary of G2L4 RT, MMEJ substrates properties and potential pathways to diverse repaired DNA products.

Most protein-DNA complexes observed in our HS-AFM datasets are composed of G2L4 RT dimer, in agreement with our previous model.^14,15^ The dissociation of G2L4 RT dimer into monomers in the presence of Mn^2+^ (**Fig. 1C**) suggests that this reverse transcriptase may act as a monomer with some functional activities on DNA substrates.^10,14^ Such Mn^2+^-induced conformational change have similarly been reported for other DNA polymerases^33^ and DNA-binding proteins^34^. Although we were unable to capture monomer-driven DSBR via MMEJ in real-time, several monomer-DNA interactions are suggestive of repair-related activities. For example. For example, the monomer can slide along the DNA backbone, possibly to search for repair sites (**Fig. 3E**), or stabilize the MH formation (**Fig. S9**).

The designed MMEJ substrates, as expected, can form MMEJ dimers via MH-mediated end joining formation; however, the four bp MH is relatively weak, with formation and dissociation dynamics observed in real-time (**Fig. 2E**). The G2L4 RT dimers preferentially bind to MMEJ substrates with a ss-overhang at the 3’ end of designed D2 strand, but they also bind to MMEJ dimers near the MH region and/or the dsDNA end (**Fig. 3A**). Upon closely inspection, we found that G2L4 RT dimers bound to DNA substrates exhibit a small topological protrusion (**Fig. 3B** and **Fig. S6**), aligning with the extruded RT3a plug reported previously.^15^ Moreover, G2L4 RT dimers can stabilize the MMEJ dimer after binding near the MH region or one of the ss-gaps, with repositioning dynamics observed in real-time (**Fig. 3C**). This behavior is consistent with hypotheses proposed in our earlier studies.^15^ With dNTPs, the MMEJ dimer can be repaired by G2L4 RT, leaving two nicked sites that require DNA ligation to complete repair (**Fig. 3H**). Notably, MMEJ dimers often adopt a remarkable bending geometry before final ligation by T4 DNA ligase (**Fig. 2D** and **Fig. 3H**), which reflects the distinct mechanical properties of dsDNA and ssDNA: dsDNA is relatively rigid and linear-like, whereas ssDNA is highly flexible and readily bent.^23,35^

Since MMEJ is an error-prone DSBR process, increasing the relative concentration of G2L4 RT compared to MMEJ sample and incubation time for the DNA repair reaction resulted in long DNA products, including linear and branched forms (**Fig. 4**). These error-prone DNA products are not easily detected by traditional molecular-biology assays due to their low concentration. Specifically, in the presence of Mn^2+^, G2L4 RT can act as a polymerase to fill ss-DNA gaps on DNA substrates with dNTPs (**Fig. 3**), but it also has terminal transferase activity that can extend the ssDNA end in a template-independent manner (**Fig. 4G** and **Fig S11**). We captured that G2L4 RT dimers can hold two MMEJ dimers by bridging their central MH regions (**Fig. 3G**), stabilize a flap ssDNA branch (**Fig. 4E-(ii)**), and promote the self-rearrangements of the DNA backbone that yield branched morphology (**Fig. 4H**). Without the DNA ligase to seal nick sites, G2L4 RT-repaired DNA molecules can undergo further elongation and backbone rearrangement to form more complex structures. Thus, in the absence of immediate ligation, increasing the enzyme-to-MMEJ ratio can drive more off-pathway elongation and branching, expanding product diversity beyond the expected MMEJ outcome. Importantly, our experimental conditions use relatively low enzyme concentrations and are therefore more likely to reflect the in vivo mechanism.

Finally, we found that addition of T4 DNA ligase largely eliminated branched DNA species, likely because ligation increases the stability of repair products (**Fig. 5**). We also captured T4 DNA ligase searching for and binding near a DNA nick via its DBD domain, demonstrating the feasibility of observing real-time DNA ligation dynamics. Recruitment of T4 DNA ligase further seals the nicked backbone, stabilizing the final DSBR product via the MMEJ mechanism and suppressing off-pathway reactions. Overall, our results not only provide new mechanistic insights into G2L4 RT-mediated MMEJ repair but also highlight the potential of HS-AFM to elucidate the dynamic interplay between polymerases and ligases during DSBR at the single-molecule level.

## ASSOCIATED CONTENT

### Supporting Information

Materials and methods, supporting figures and tables, and supporting movie captions.

## AUTHOR INFORMATION

## Author contributions

P. Z., M. G., Y.J.Z., and Y.-C.L. designed research. M. G. and Y.J.Z. designed and prepared the DNA samples. P. Z. performed HS-AFM experiments. P. Z. and Y.-C.L. analyzed the data and wrote the manuscript. Y.J.Z. and A.M.L. reviewed and edited the manuscript, including relating new findings to previous studies of G2L4 RT.

## Acknowledgements

This work was supported by start-up fund from the University of Texas at Austin and the U.S. National Institutes of Health grant R35 GM150528 to P. Z. and Y.-C. L.; R35 GM148356 and R01CA281106 to Y.J.Z.. Research on this topic in the A.M.L. laboratory was supported by US National Institutes of Health grant R35 GM136216 and Welch Foundation grant F-1607.

## Data availability

The manuscript figures, supplementary figures, and supplementary movies contain all data necessary to interpret, verify and extend the presented work. The raw data files can be obtained from the authors upon reasonable request.

