## Supplemental Information for "Real-Time Visualization of G2L4 Reverse Transcriptase in DNA Repair via Microhomology-Mediated End Joining"

### This PDF includes:

***Materials and Methods*** ***2***
G2L4 RT Expression and Purification 2
Factor Xa Cleavage of the N-terminal MBP Tag 2
MMEJ Substrate Preparation 2
HS-AFM Imaging 2
Image Processing and Data Analysis 2
***Supplementary Tables*** ***4***
Table S1. MMEJ substrate sequence 4
Table S2. Experimental conditions of HS-AFM experiments 4
***Supplementary Figures*** ***5***
Figure S1. Structural dynamics of a G2L4 RT dimer in APO condition. 5
Figure S2. Relative percentage of MMEJ dimers observed under different conditions. 5
Figure S3. Regional characterization of MMEJ DNA sample. 6
Figure S4. Representative HS-AFM images of MMEJ sample observed in the presence of 1 mM dNTPs and 10 mM $Mg^{2+}$. 7
Figure S5. Time-lapse HS-AFM images of a G2L4 RT dimer clamping at MH of a MMEJ dimer. 7
Figure S6. G2L4 RT dimer bound to MMEJ substrate with a topological extrusion of RT3a domain. 8
Figure S7. Representative HS-AFM images of G2L4 RT dimer with different number of RT3a plug protrusion. 9
Figure S8. Representative HS-AFM image of G2L4 RT pre-incubated with MMEJ sample with dNTPs and $Mn^{2+}$ for one hour. 9
Figure S9. G2L4 monomer can bridge MH formation in the presence of $Mn^{2+}$. 10
Figure S10. Structure characterization of MMEJ dimers under different conditions. 10
Figure S11. Terminal transferase activity of G2L4 RT in the presence of $Mn^{2+}$ and dNTPs. 11
***Description of Supplementary Movies*** ***12***

### *Materials and Methods*

#### G2L4 RT Expression and Purification

N-terminal MBP-tagged G2L4 RT was expressed and purified as described.[1] Protein purity is polished by SEC and checked by SDS-PAGE. Protein is concentrated to 8-10mg/ml in 20 mM Tris-HCl pH 7.5, 50 mM NaCl and 10% glycerol and stored at -80℃ until used.

#### Factor Xa Cleavage of the N-terminal MBP Tag

To prepare G2L4 RT for HS-AFM experiments, the N-terminal MBP tag was removed by Factor Xa as previously described.[1] Following cleavage, the protein sample was purified using MBP affinity column and checked by SDS-PAGE. The protein was diluted to 1-2 mg/mL in buffer containing 20 mM Tris-HCl pH 7.5, 50 mM NaCl and 10% glycerol at 4℃ and stored at -80℃ until used.

#### MMEJ Substrate Preparation

MMEJ substrate was prepared by annealing a 96-nt DNA oligonucleotide (Top) ending with a 4-nt microhomology (CCGG) to a complementary 60-nt DNA oligonucleotide with a 3' inverted dT blocking group (Bottom) leaving a 36-nt single-stranded 3' overhang. To anneal the complementary strands of the MMEJ substrate, 10 µM Top and 20 µM Bottom oligonucleotides in 100 µL TE buffer were heated to 95°C for 3 min and cooled at 0.1°C/min to 25°C in a T100 thermal cycler (Bio-Rad). Both DNA oligonucleotides were purchased from Integrated DNA Technologies (IDT) and DNA sequences are listed in **Table S1**.

#### HS-AFM Imaging

All HS-AFM movies were taken by amplitude-modulation mode HS-AFM (RIBM, Japan) and imaged in the buffer solutions. Ultrashort Cantilevers (NanoWorld) with a spring constant of ~0.15 N/m, resonance frequency of ~600 kHz, and a quality factor of ~1.5 in buffer solutions were used.

Experimental conditions of HS-AFM experiments are summarized in **Table S2**. The imaging buffer for all experiments contained 20 mM Tris-HCl (pH 7.5), 50 mM NaCl and 10 mM $MgCl_2$, supplemented with $MnCl_2$, dNTPs and ATP as specified in **Table S2**. To deposit the sample on substrate for imaging, the freshly cleaved mica was pre-incubated with 0.05% (v/v) APTES, and then gently rinsed with Milli-Q water. A total of 1-2 µL of premixed sample was diluted to proper concentration and deposited on APTES-treated mica surface for 5 mins, gently rinsed with the imaging buffer prior to HS-AFM experiments.

#### Image Processing and Data Analysis

All HS-AFM images were processed in ImageJ with drift correction, background flattening, scanning noise removal, and contrast adjustment. The volume of G2L4 RT particles was analyzed using Gwyddion by properly thresholding the HS-AFM images and measuring grains properties. Simulated AFM images (4 pixels/nm resolution) were generated using Mat-SimAFM[2] by convolving the protein's PDB structure by a cone-shape tip (tip radius 1 nm, cone angle 15°). Kymographs were generated in ImageJ by reslicing the HS-AFM movies along the DNA backbone, followed by a maximum intensity Z-projection to visualize protein position over time. Intermolecular distances between G2L4 monomers were calculated from the x, y coordinates of protein centers. Protein particles were detected using Laplacian of Gaussian (LoG) detector, and the center coordinates were extracted using Trackmate plugin in Fiji.[3] Finally, all data analysis were plotted using custom-written Python scripts.

### *Supplementary Tables*

#### Table S1. MMEJ substrate sequence

| Name | Sequence |
|---|---|
| 96-nt DNA<br>4 microhomology (CCGG)<br>Top | 5'CCCTGTACAGTAAGAGCCTACTCATGGATCCTCCTTGTG<br>ATGTAAGGTACAGTAAGATGTGCCTACTCATGGTCGGATC<br>CTCCTTGTGATGCCCGG 3' |
| 60-nt DNA 3' block<br>(Inverted dT at the 3'-end)<br>Bottom | 5'ACATCTTACTGTACCTTACATCACAAGGAGGATCCATGA<br>GTAGGCTCTTACTGTACAGGG/ InvdT / 3' |

#### Table S2. Experimental conditions of HS-AFM experiments

| Figures | System | Molecular Players | | | Imaging Buffer | | | | Mixing Molecular Players | |
|---|---|---|---|---|---|---|---|---|---|---|
| | | G2L4 | MMEJ | T4 ligase | $Mg^{2+}$ | $Mn^{2+}$ | dNTPs | ATP | Pre-incubation | Live addition |
| 1B(i),1C(i),1D, S1 | G2L4-APO | 50 nM | | | 10 mM | | | | | |
| 1B(ii), 1C(ii) | G2L4-$Mn^{2+}$ | 50 nM | | | 10 mM | 5 mM | | | | |
| 2, S2, S3, S4 | MMEJ | | 10 nM | | 10 mM | | | | | |
| 3A, 3B, S5, S6, S7 | G2L4+MMEJ | 500 nM | 50 nM | | 10 mM | | | | 30 mins~ 2 h | |
| 3C, 3D, 3G, S8, S9 | G2L4 repair | 500 nM | 50 nM | | 10 mM | 1 mM | 1 mM | | | √ |
| 3E, 3F, 3H | G2L4 repair | 500 nM | 10 nM | | 10 mM | 1 mM | 1 mM | | 6 h | |
| 4A | G2L4 repair-molar ratio 10:1 | 100 nM | 10 nM | | 10 mM | 1 mM | 1 mM | | 6 h | |
| 4A | G2L4 repair-molar ratio 50:1 | 500 nM | 10 nM | | 10 mM | 1 mM | 1 mM | | 6 h | |
| 4C, 4E, 4F, 4G, S10 | G2L4 repair-elongated and branched products | 500 nM | 10 nM | | 10 mM | 1 mM | 1 mM | | 6 h | |
| 5 | DNA ligation | 500 nM | 50 nM | 0.04 U | 10 mM | 1 mM | 1 mM | 1 mM | 6 h(repair)<br>Overnight (ligation) | |

### *Supplementary Figures*

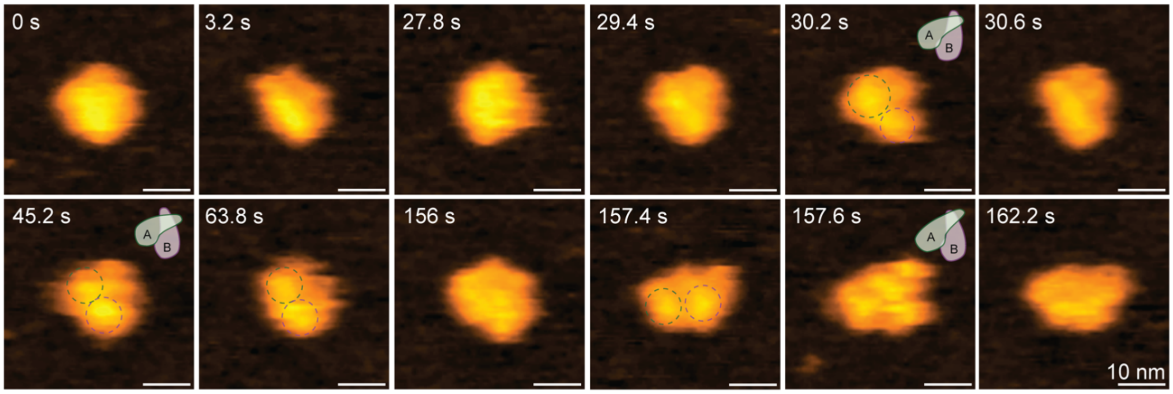


**Figure S1. Structural dynamics of a G2L4 RT dimer in APO condition.** Representative HS-AFM images of a G2L4 RT dimer show certain conformational dynamics with bi-lobed features (**Movie S2**). At 30.2 s and 45.2 s, the observed structures adopt a dimeric conformation similar to the crystal structure (**Fig. 1D**). At 30.2 s, 45.2s, 63.8 s, and 157.4 s, two palm-finger domains (bi-lobed features, dashed circles) appear slightly separated in a side-by-side arrangement. More fine structural features of G2L4 RT dimer are observed at 157.6 s.

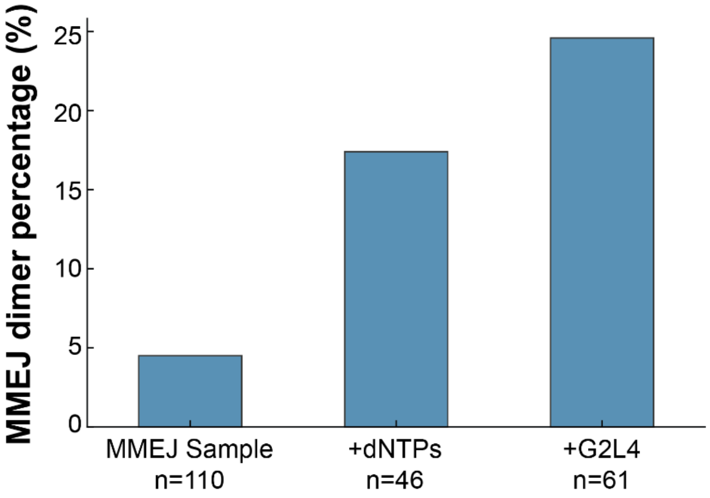


**Figure S2. Relative percentage of MMEJ dimers observed under different conditions.**

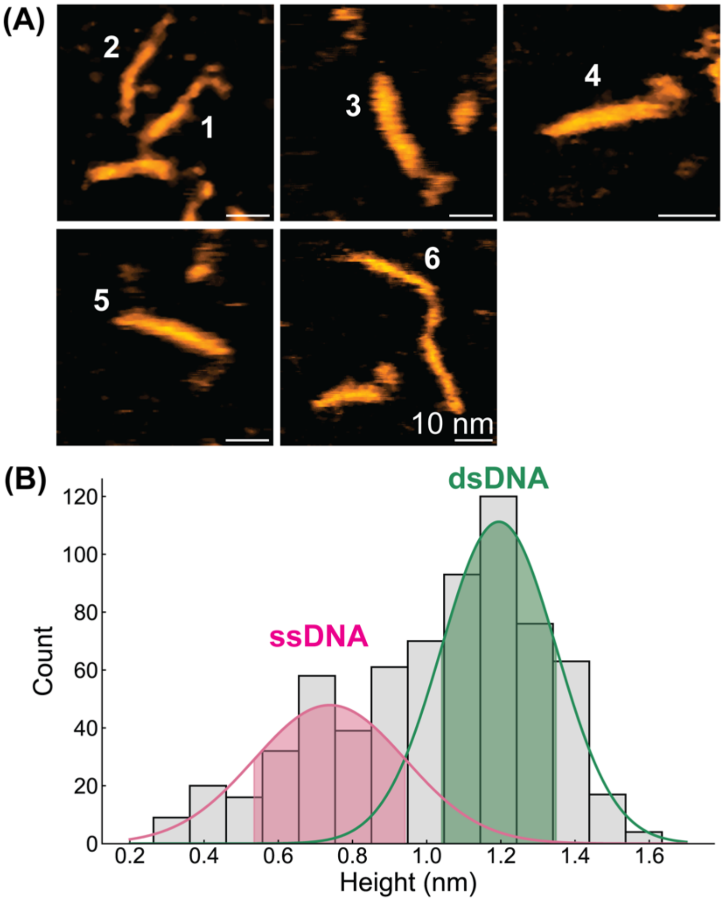


**Figure S3. Regional characterization of MMEJ DNA sample. (A)** Representative HS-AFM image of MMEJ substrates used to analyze the regions of ssDNA and dsDNA. **(B)** Accumulative height histograms of the MMEJ substrates masking with a height threshold of 0.3 nm. Two Gaussian fits centered at 0.7 ± 0.2 nm for ssDNA and 1.2 ± 0.2 nm for dsDNA. We note that the mica background is flattened with a mean value of approximately at 0 in height.

125

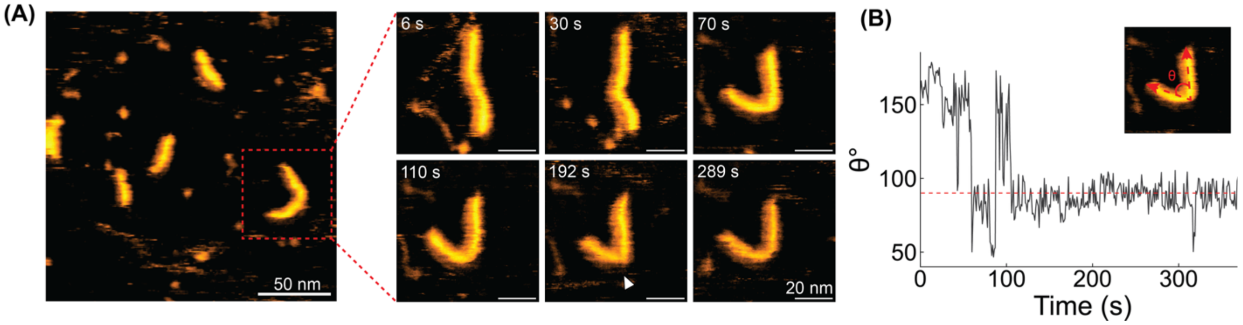


**Figure S4. Representative HS-AFM images of MMEJ sample observed in the presence of 1 mM dNTPs and 10 mM $Mg^{2+}$. (A)** Time-lapse HS-AFM images of a MMEJ dimer without dissociation. The sharp bend, indicated by white arrowhead, highlights the DSBR is incomplete purely with dNTPs filling up the ss-gaps. **(B)** Tracked bending angle between two ends of the observed MMEJ dimer. Red dashed line represents 90 degrees.

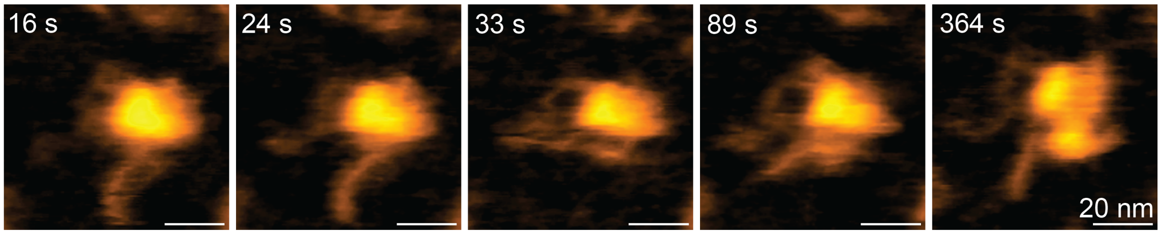


**Figure S5. Time-lapse HS-AFM images of a G2L4 RT dimer clamping at MH of a MMEJ dimer.** Although the G2L4 RT dimer shows certain conformational changes, the protein-DNA complex remains stable during the observation.

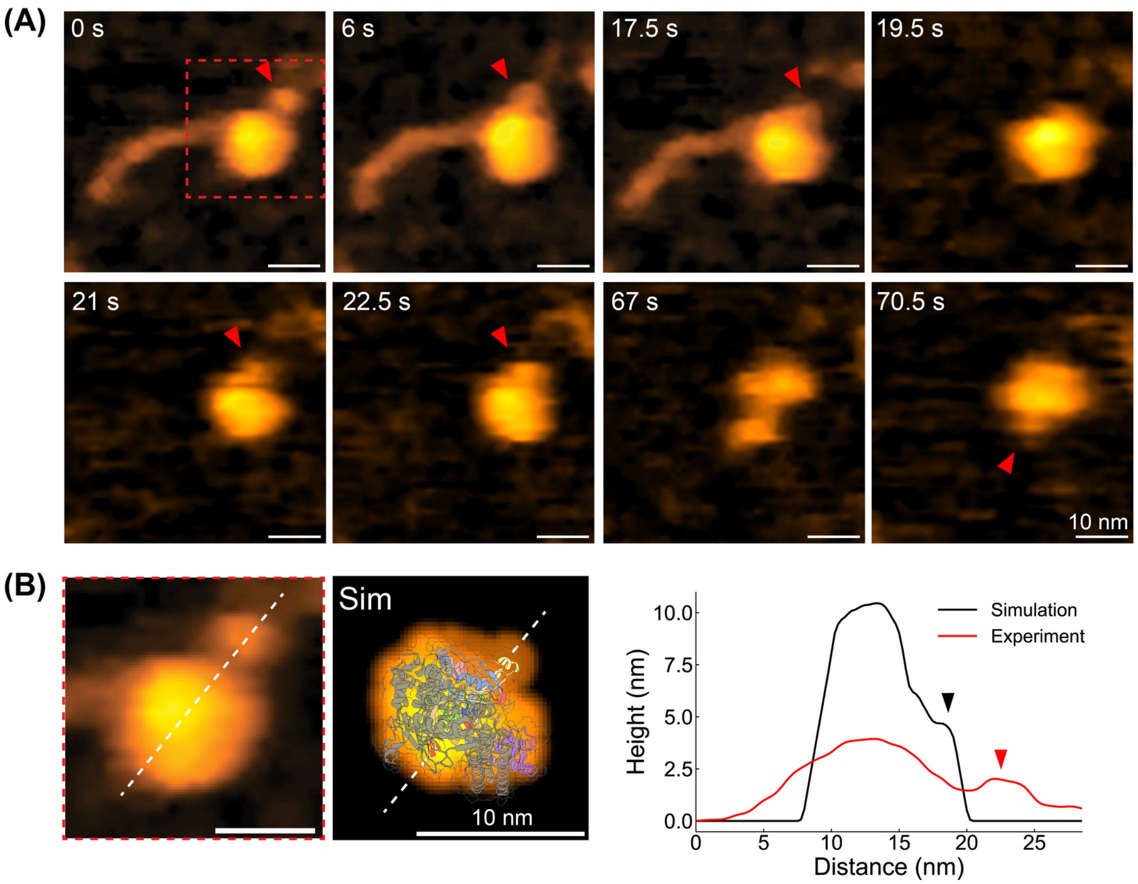


**Figure S6. G2L4 RT dimer bound to MMEJ substrate with a topological extrusion of RT3a domain. (A)** Representative HS-AFM images showing a small RT3a plug protrusion (red arrows) when G2L4 RT dimer binds to MMEJ substrate. **(B)** The comparison between zoomed-in HS-AFM image of the G2L4 RT dimer and simulated AFM topography of G2L4 RT dimer in complex with 15-nt snapback substrate (PDB: 9D4S; molecular orientation is adjusted and viewed from the side to highlight RT3a plug). Right: Height cross-sectional profile of the experimental and simulated AFM data along the white dashed lines. The arrows indicate the position of RT3a plug. The significant height difference between experimental and simulated AFM data is due to the G2L4 RT is free-standing above the background substrate in simulated AFM topography.

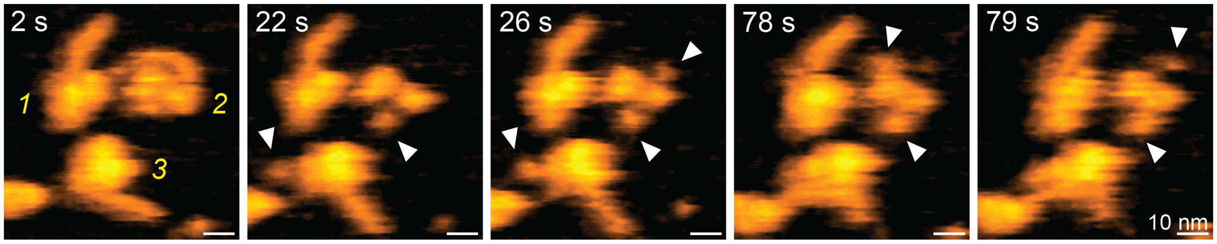


**Figure S7. Representative HS-AFM images of G2L4 RT dimer with different number of RT3a plug protrusion.** Three G2L4 RT dimers, highlighted by yellow numbers, individually bind to a MMEJ substrate (t = 2 s). For G2L4 RT#2, the MMEJ substrate dissociates from this protein with a bi-lobed feature, where two RT3a plug extrusions are observed at the opposite sides, indicated white arrows.

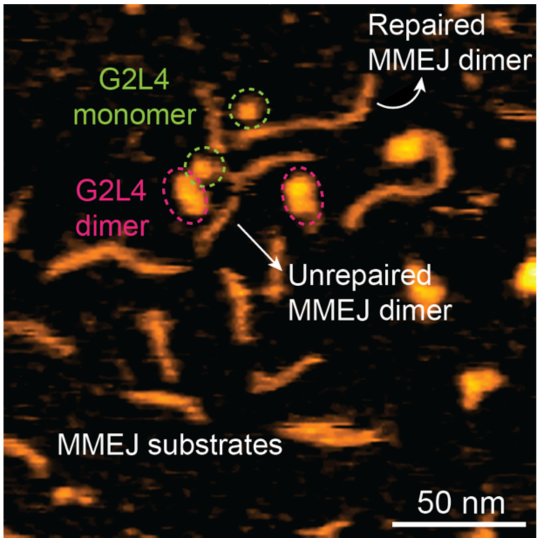


**Figure S8. Representative HS-AFM image of G2L4 RT pre-incubated with MMEJ sample with dNTPs and $Mn^{2+}$ for one hour.** G2L4 monomers, dimers, and both repaired and unrepaired MMEJ dimers are coexisted.

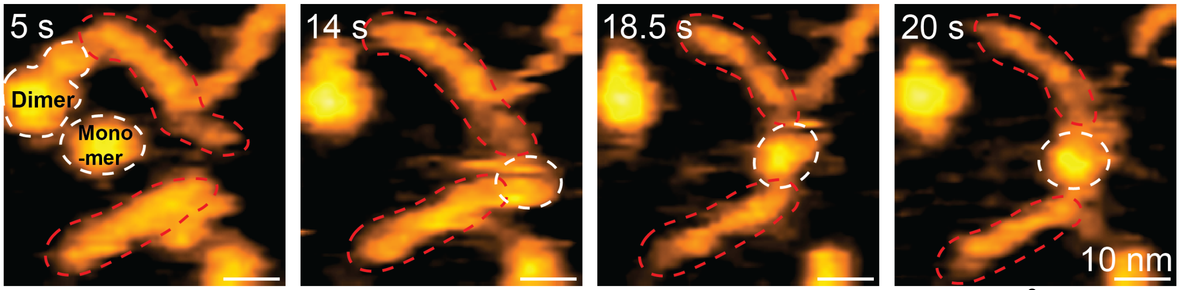


**Figure S9. G2L4 monomer can bridge MH formation in the presence of $Mn^{2+}$.** At t = 5 s, two MMEJ substrates are separated as highlighting by red dashed circles. After 14 s, a G2L4 monomer bridged two ss-DNA overhangs together, forming MH.

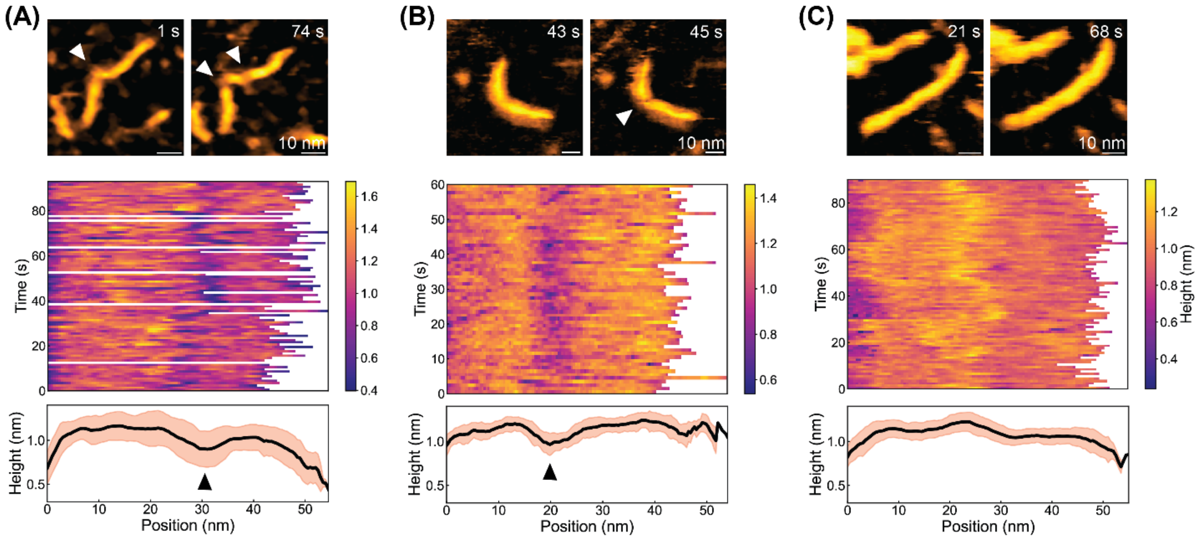


**Figure S10. Structure characterization of MMEJ dimers under different conditions.** Each panel displays (*top*) representative HS-AFM images, (*middle*) kymographs of height along the DNA backbone, and (*bottom*) average height profile (solid black line) with standard deviation, orange shading, for (**A**) MMEJ sample alone, **(B)** MMEJ sample incubated with dNTPs, and **(C)** MMEJ dimer incubated with G2L4 RT, dNTPs and $Mn^{2+}$.

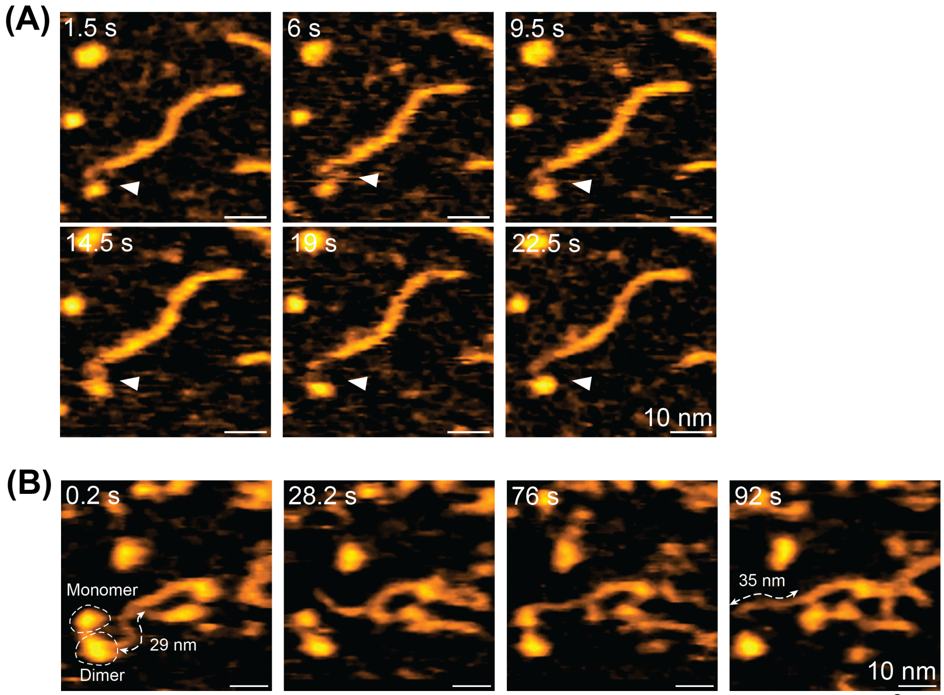


**Figure S11. Terminal transferase activity of G2L4 RT in the presence of $Mn^{2+}$ and dNTPs. (A)** Time-lapse HS-AFM images of a G2L4 RT monomer interacting with ssDNA overhang, indicated by white arrowheads. No significant elongation is observed in the trajectory. **(B)** Realtime ssDNA elongation by G2L4 RT molecules. The ssDNA length is measured by setting high threshold at 0.8 nm, and increased by ~ 6 nm (~30 nt; by assuming 0.2 nm/nt obtained in **Fig. 2C**) at 92 s.

143

***Description of Supplementary Movies***

**Supplementary Movie S1. Structural dynamics of APO-G2L4 RT dimer.** HS-AFM movie of an APO-G2L4 RT dimer deposited on freshly cleaved mica (see **Fig. 1C-(i)**). The G2L4 RT molecule was aligned at the center for enhancing the conformational dynamics. Imaging parameters: 2 frames/s, 0.25 nm/pixel.

**Supplementary Movie S2. Structural dynamics of another APO-G2L4 RT dimer.** HS-AFM movie of another APO-G2L4 RT dimer deposited on freshly cleaved mica (see **Fig. 1D** and **Fig. S1**). Imaging parameters: 5 frames/s, 0.5 nm/pixel.

**Supplementary Movie S3. Dissociation of a G2L4 dimer in $Mn^{2+}$ condition.** HS-AFM movie of a G2L4 RT dimer observed in buffer containing 6 mM $Mn^{2+}$ (see **Fig. 1C-(ii)**). At the beginning of the movie (~1 s), a monomer and a dimer coexisted in the imaging area. The dimer was intact at ~13 s and then began showing conformational changes with two lobed-like structure occasionally visible. Two monomers fully dissociated at ~182 s. Imaging parameters: 1 frame/s, 0.375 nm/pixel.

**Supplementary Movie S4. Structural dynamics of individual MMEJ substrates.** HS-AFM movie of three individual MMEJ substrates deposited on APTES-coated mica (see **Fig. 2B**). The MMEJ substrates, especially the middle one, showed clear height difference along the DNA backbone, indicating the regions of dsDNA and ssDNA. ssDNA overhang was more flexible compared to dsDNA end. Imaging parameters: 1 frame/s, 0.5 nm/pixel.

**Supplementary Movie S5. Structural dynamics of a MMEJ dimer.** HS-AFM movie of a MMEJ dimer deposited on APTES-pretreated mica (see **Fig. 2D**). The MH region and ss-gap could be clearly identified by the height variation along the backbone. The dsDNA region was freely rotating as an arm relative to the ss-gap as a hinge. Imaging parameters: 1 frame/s, 0.5 nm/pixel.

**Supplementary Movie S6. Association and dissociation of a MMEJ dimer.** HS-AFM movie of realtime association and dissociation of a MMEJ dimer (see **Fig. 2E**). In this scanning area, there initially were a MMEJ substrate and a transiently formed MMEJ dimer. In the absence of G2L4 RT, the MH dissociated at ~3.5 s, associated again at ~4 s, and finally separated after 6 s. Imaging parameters: 2 frames/s, 0.286 nm/pixel.

**Supplementary Movie S7. A MMEJ dimer clamped by a G2L4 RT dimer.** HS-AFM movie of a G2L4 dimer clamping at the MH region between two MMEJ substrates (see **Fig. S5**). Imaging parameters: 1 frame/s, 0.5 nm/pixel.

**Supplementary Movie S8. G2L4 RT dimer with a RT3a plug protrusion upon MMEJ substrate engagement.** HS-AFM movie showing a small RT3a plug protrusion when G2L4 RT dimer binds to MMEJ substrate (see **Fig. 3B** & **Fig. S6**). Imaging parameters: 2 frames/s, 1 nm/pixel.

**Supplementary Movie S9. G2L4 RT dimer repositions on a MMEJ dimer.** HS-AFM movie of a G2L4 RT dimer repositioning on a MMEJ dimer (see **Fig. 3C**). The imaging was performed on 0.05% APTES coated mica, with repair reaction mixture added realtime into liquid cell. The G2L4 RT dimer was firstly bound to the right ss-gap, repositioned to the left ss-gap (~37 s), bounced back and forth twice, and finally bound to the left ss-gap. At 94 s, another G2L4 RT dimer bound to the left side of DNA. Imaging parameters: 2 frames/s, 1 nm/pixel.

**Supplementary Movie S10. Molecular interactions between G2L4 RT and long DNA products.** HS-AFM movie of an intermediate DNA product interacting with several G2L4 RT molecules (see **Fig. 4E**). Along the DNA backbone, three events were happening simultaneously: G2L4 dimer sliding, unidirectional repair and strand annealing. Imaging parameters: 1 frame/s, 1.5 nm/pixel.

**Supplementary Movie S11. Self-rearrangements of the DNA backbone at a nicked site to form a branched morphology.** HS-AFM movie of an intermediate DNA product showing strand rearrangement (see **Fig. 4H**). The long linear DNA product was broken at a nicked site (~93 s). The broken end rearranged to the middle of DNA backbone, forming a branched structure. Imaging parameters: 1 frame/s, 1 nm/pixel.

**Supplementary Movie S12. T4 DNA ligase repairs a MMEJ dimer**. HS-AFM movie of a T4 DNA ligase interacting with a MMEJ dimer with a nick site (see Fig. **5D-5E**). A T4 DNA ligase in solution appeared to bind to the MMEJ dimer (~23 s). The DBD domain initiated the DNA engagement, slid to search nick site (23~27 s). After transient domain rearrangement (28~29 s), the DNA backbone is ligated after 38 s. Imaging parameters: 1 frame/s, 0.25 nm/pixel.
